# Distribution and diet of Central American Clouded Tiger Cat *Leopardus pardinoides oncilla* via noninvasive genetics

**DOI:** 10.64898/2026.05.26.727971

**Authors:** Torrey W. Rodgers, Roberto Salom-Pérez, Stephanny Arroyo-Arce, Erick Viquez-Alvarado, Pedro L. Castillo-Caballero, Daniela Araya-Gamboa, Claudio M. Monteza-Moreno, Michael S. Mooring, Marta Vargas, Karen E. Mock

## Abstract

The Clouded Tiger Cat *Leopardus pardinoides* is a recently recognized Neotropical species for which ecological and natural history data are sparse. Knowledge of species distribution and elevational range are largely based upon camera trap studies, and its diet has not been examined. The objective of this study was to better define the geographical and elevational distribution of the subspecies *L. pardinoides oncilla* in Central America using genetically confirmed records. We also provide the first diet analysis for *L. pardinoides.* We conducted extensive surveys for scat samples by visual means and with a scent detection dog across mountain ranges in Panama and Costa Rica, confirmed species identity using Sanger sequencing of mitochondrial DNA, and analyzed diet composition via DNA metabarcoding. Altogether, we collected 195 confirmed *L. pardinoides* scats. Records were strongly associated with high elevations, with a median elevation of 2805 m, and a maximum of 3422 m. Our findings highlight core habitat for *L. pardinoides* in the Cordillera Talamanca mountains in the border region of Panama and Costa Rica, and we identified isolated populations from the Central Volcanic Cordillera of Costa Rica and the Central Cordillera of Panama. DNA metabarcoding detected 59 vertebrate taxa. Diet was dominated by small mammals (74% of samples), especially cricetid rodents and shrews. Birds were also commonly detected (35% of samples), while reptiles (13%) and amphibians (1.9%) were less frequent. Estimated median adult prey mass was low (25 g), indicating that the species specializes on small prey. Most vertebrate prey identified were species endemic to montane regions of Panama and Costa Rica. These findings provide valuable ecological and natural history data on Clouded Tiger Cats in Central America that will inform conservation and management of this recently recognized species.

## Introduction

Felid mesopredators such as those from the genus *Leopardus* play important roles in Neotropical ecosystems (de Oliveira and Pereira 2014) however, many remain poorly studied compared to their larger counterparts (Gálvez *et al*. 2023). Basic natural history knowledge about distribution and diet is foundational for conservation and management of these species (Greene 2005), as well as for improved understanding of their ecological role in Neotropical mammal community dynamics. One group of Neotropical felids, the Tiger Cat or *Leopardus tigrinus* complex, has a history of taxonomic and ecological confusion (Giordano 2012; Bonilla-Sánchez *et al*. 2024). This species complex was once considered to be a single species, *Leopardus tigrinus,* with a broad geographic and elevational distribution covering much of tropical South America and part of Central America. Recent research has provided strong evidence based on morphology (de Oliveira *et al*. 2024), genetics (Lescroart *et al*. 2023), and ecology and biogeography (Bonilla-Sánchez *et al*. 2024; de Oliveira *et al*. 2024) that this complex is composed of at least three distinct species: *L. tigrinus, L. guttulus* (Trigo *et al*. 2013a), and the newly recognized *L. pardinoides* (de Oliveira *et al*. 2024).

The most recently recognized species from this complex, the Clouded Tiger Cat, *Leopardus pardinoides* (de Oliveira *et al*. 2024), occurs in the Andean region of South America, the Talamanca mountain of Panama and Costa Rica, and the central volcanic cordillera of Costa Rica. There are no known continuous populations between the Andes and Talamanca, hence, disjunct Central American populations are considered a subspecies *L. p. oncilla*, often referred to by the common name Oncilla (Ramírez-Fernández *et al*. 2024). Recent *L. p. oncilla* research with camera trapping has determined that the species is associated with rugged high elevation terrain and is most commonly found in montane cloud forest (de Oliveira *et al*. 2024; Ramírez-Fernández *et al*. 2024). This subspecies currently faces threats from habitat loss and fragmentation, particularly from conversion of forest habitat to agriculture (de Oliveira *et al*. 2024). Other threats include isolation of populations, which will likely be exacerbated by climate change-caused upward contraction of montane cloud forest and páramo habitat, resulting in loss of high elevation refugia (Ramírez-Fernández *et al*. 2024).

Although camera trap studies have expanded knowledge of Clouded Tiger Cat occurrence with model-predicted distributions (de Oliveira *et al*. 2024; Ramírez-Fernández *et al*. 2024), some uncertainties remain. For instance, elevational limits and use of páramo habitat at high elevations above montane cloud forests are unclear. Moreover, the true eastern and western range limits for the subspecies remain uncertain, especially in isolated mountain ranges with suitable habitat but poor connectivity with core habitat in the Talamanca mountains. These isolated mountain ranges could function as high elevation refugia for the subspecies, although isolation could pose conservation challenges. Camera traps have limitations for studying small elusive felids because detections can only be captured at point locations where cameras are placed. This constrains detections from high-elevation páramo habitats where camera trap placement and maintenance is more challenging due to accessibility and habitat openness, with lack of game trails and trees for camera placement. Additionally, although experts can identify *L. pardinoides* from camera trap photos with relatively high accuracy (de Oliveira *et al*. 2024), distinguishing *L. pardinoides* from Margay (*L. wiedii)* or juvenile Ocelots from camera trap images can be challenging (Anderson *et al*. 2024), potentially leading to false presences.

The diet of *Leopardus pardinoides* is unknown from anywhere within the species range. Diet has been studied in the closely related species *Leopardus tigrinus* based on microscopic analysis of prey remains from scat (Silva-Pereira et al., 2011; Wang, 2002), and roadkill stomach contents (Trigo et al., 2013), in Brazil. Despite limitations from small sample sizes (36 scat samples, Silva-Pereira et al., 2011; 24 scat samples, Wang, 2002; 13 stomach contents, Trigo et al., 2013) and the limitation of identifying most prey items beyond family or genus based on morphology, these studies found that small mammals were the most important prey, followed by birds. All three studies reported infrequent reptile prey; one study reported a single occurrence of an unidentified reptile, another reported a single occurrence of an unidentified snake, and the third study reported four occurrences of snake remains identified to family Colubridae. Amphibians were not identified as prey items in any of these *L. tigrinus* studies. Likewise, traditional diet studies relying on morphological identification of prey remains can pose limitations. Traditional methods require trained expertise in identifying prey from bones and hair and often provides only coarse taxonomic resolution to family or genus level (de Sousa *et al*. 2019). Traditional diet methods may also miss soft-bodied prey such as small or juvenile birds and small mammals, as well as reptiles and amphibians (Thuo *et al*. 2019).

We hypothesized that the diet of *L. pardinoides* may differ from *L. tigrinus* due to several factors. First, although *L. pardinoides* was until recently recognized as the same species as *L. tigrinus*, they are now recognized as separate species occupying a very different ecoregion (Bonilla-Sánchez *et al*. 2024; de Oliveira *et al*. 2024). All previous diet studies of *L. tigrinus* were conducted in the lowland forest or grassland ecosystems of Brazil where *L. tigrinus* is present, habitat that is strikingly different from montane cloud forest ecosystems of the Andes and Central America that have more vertical structure and greater endemism. Thus, in contrast to *L. tigrinus*, we predicted that *L. pardinoides* may consume more arboreal prey and more birds, as well as small mammals, birds, reptiles, and amphibians endemic to montane cloud forests.

In this study, we combine non-invasive genetic methods from scat sampling (Rodgers and Janečka 2013) with DNA metabarcoding (Taberlet *et al*. 2012) to provide key knowledge on the ecology and natural history of the Clouded Tiger Cat in Central America. First, we genetically confirm the distribution and elevational range of *L. p. oncilla* in Panama and Costa Rica. We provide records from many new locations, and we identify core areas of species occurrence and regions where Clouded Tiger Cat conservation actions and fine scale research may be most needed. Second, we provide the first dietary analysis for *L. pardinoides and* evaluate the reliance of *L. p. oncilla* on endemic cloud forest prey. Together, these results contribute substantially to knowledge of Clouded Tiger Cat ecology and behavioral and provide data on prey base essential for conservation planning.

## Materials and methods

### Field methods

To determine the distribution of *L. p. oncilla* in Central America, we used noninvasive genetic methods from scat samples (Rodgers and Janečka 2013). Scat surveys were conducted between Guanacaste Province, the westernmost province of Costa Rica, to Darien Province, the easternmost province in Panama (Table 1). Scat samples were collected from both Panama and Costa Rica utilizing visual searches by human investigators, and from Costa Rica using a detection dog to locate scats by scent (McKeague *et al*. 2024). A detection dog was not used in Panama because dogs are not permitted in Panamanian protected areas. For visual searches, we collected scat samples primarily along trails, and occasionally off trail when conditions permitted. We conducted a total of 74 field-days of visual searches, 31 days in Costa Rica and 43 days in Panama, between 2020-2025 (Table 1). Visual sampling was typically conducted in remote areas by a team of two people, although in some instances (e.g., longer expeditions) the team consisted of 3-5 people, and in rare instances sampling was conducted by a single individual.

**Table 1.**
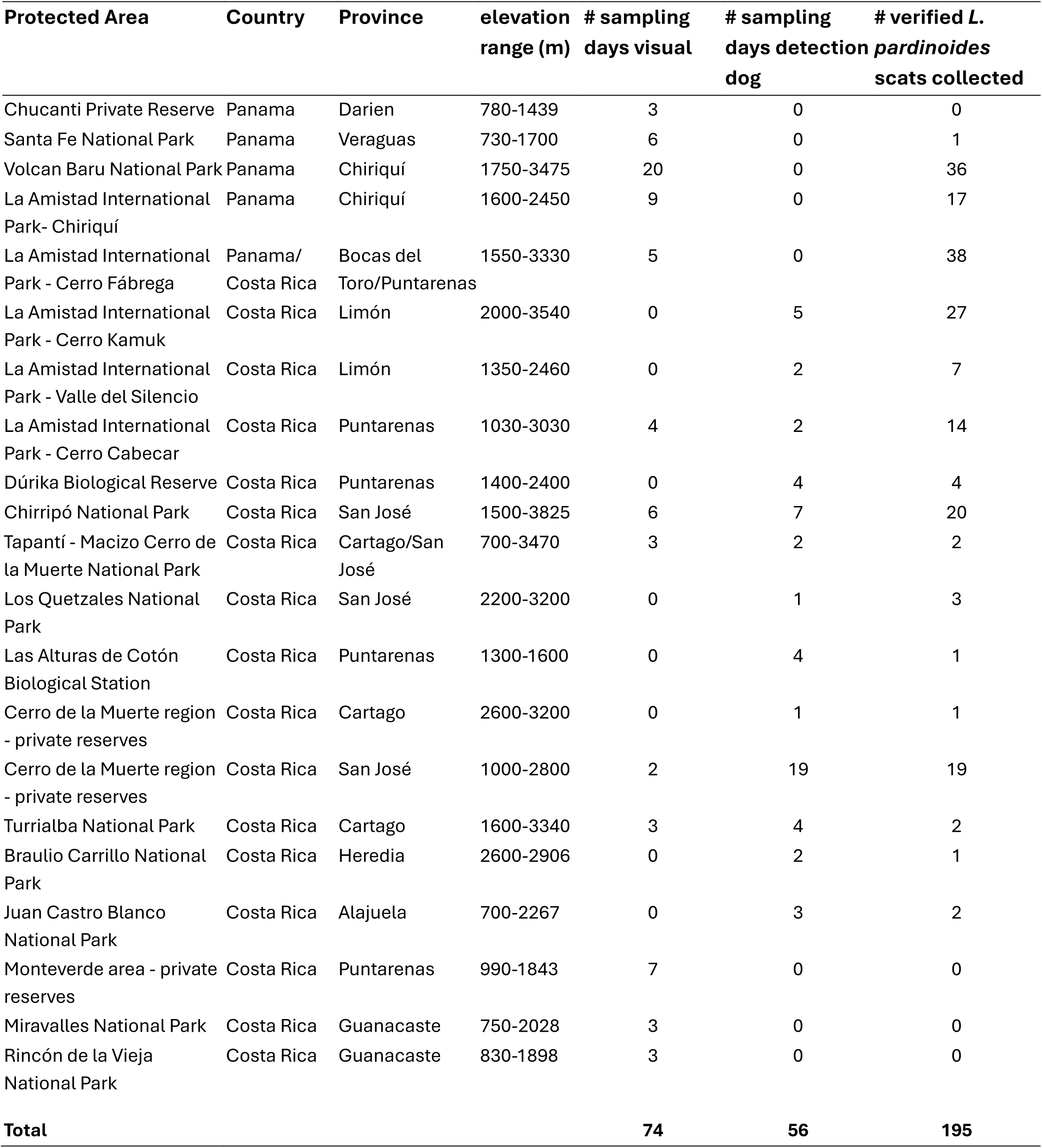
Protected areas sampled and verified *Leopardus pardinoides* scats collected in Panama and Costa Rica by visual and detection dog sampling.

We conducted detection dog sampling in Costa Rica from potential *L. p. oncilla* habitat (habitat > 1000 m elevation) for a total of 56 days between 2013-2023 (Table 1). We also conducted 81 days of detection dog sampling in habitats below 1000 m (Supplementary Table S1). The detection dog team consisted of a trained dog, its handler, and a field assistant. Surveys used a male yellow Labrador Retriever named Tigre (2019-2023) and a male German Shorthaired Pointer named Google (2013-2016), trained to detect scats from all six wild cat species inhabiting Costa Rica (*Panthera onca, Puma concolor, Herpailurus yagouaroundi, L. pardalis, L. wiedii, and L. pardinoides*). At each site, searches focused on roads, trails, off-trail areas, and streams. The handler and dog walked ahead of the team; the dog worked off leash and was trained to sit when a felid scat was detected, followed by a reward.

When a scat was found, we removed a 2-3 cm^3^ piece of scat material and placed it in a 15 ml falcon tube filled ¾ full of silica gel, with a kimwipe separating the sample from the silica (for visual sampling) or in a 50 ml plastic jar filled with silica gel (for detection dog sampling). Samples were stored at room temperature for 1-2 weeks during sampling, after which they were transported to a -20° C freezer at Panthera Costa Rica headquarters in San Jose, Costa Rica, or the Smithsonian Tropical Research Institute in Panama City, Panama. Each year, after obtaining necessary export permits, samples were shipped to the Molecular Ecology Laboratory at Utah State University, where they were stored in a -80° C freezer until we conducted laboratory work.

### Laboratory methods

We performed scat extractions with a QIAamp Fast DNA Stool Mini kit (QIAGEN, Hilden, Germany). We cut an approximately 0.5 to 1 cm^3^ portion of scat into a sterile petri dish with sterile scissors and added 2 ml of QIAGEN inhibitEX Buffer. We chopped up the sample with sterile scissors and forceps until it was fully suspended in the buffer and then moved 1 ml of buffer with suspended scat material to a 1.7 ml centrifuge tube. We then proceeded with the extraction following the manufacturer’s protocol for stool samples for DNA analysis. Each round of extractions included a blank negative control with buffer that went through all the same steps as scat samples to monitor for contamination.

Species ID of scat samples was determined with Sanger sequencing using the primers ATP6-DF3 (Chaves *et al*. 2012) and Ltio-ATP6-R (Rodgers and Kapheim 2017). These primers, which were modified specifically for amplification of DNA from *L. p. oncilla* (Rodgers and Kapheim 2017), target a 172 base pair (including primers) fragment of the mitochondrial gene ATP6, which is reliable for distinguishing Neotropical carnivore species (Chaves *et al*. 2012). PCR reactions contained 12.5 μl of DreamTaq Hot Start PCR Master Mix (Thermo Fisher Scientific, Waltham, Massachusetts, USA), 0.4 uM forward and reverse primers, 0.05 μg of bovine serum albumin and 4 μl of scat extraction template in a total reaction volume of 25 μl. PCR cycling conditions were: initial denaturation at 95° C for 3 min, 50 cycles of 95° C for 30 sec, 54° C for 30 sec, and 72° C for 60 sec, followed by a final elongation of 72° C for 7 minutes. All PCRs included no-template negative controls, and we also conducted PCR on extraction negative controls to monitor for contamination. We obtained species assignments from sequence data with the online tool DNA Surveillance Carnivora (https://dna-surveillance.auckland.ac.nz).

For diet metabarcoding, we conducted a two-stage library preparation protocol for Illumina sequencing on 190 of the 195 total samples confirmed as *L. pardinoides* by Sanger sequencing, with the same extractions used for species ID. We conducted an initial PCR with the primers mlCOIintF (5’-GGWACWGGWTGAACWGTWTAYCCYCC-3’; Leray et al., 2013) and primer jgHCO2198 (5’-TAIACYTCIGGRTGICCRAARAAYCA-3’; Geller et al., 2013). PCR reactions contained10 μl of Amplitaq Gold 360 Master Mix (Thermo Fisher Scientific), 0.5 uM of each of forward and reverse primer containing Illumina flow cell sequencing adapters, 0.04 μg bovine serum albumin, and 4 μl of extraction template in a total reaction volume of 20 μl. PCR conditions were: initial denaturation of 10 min at 95° C, 35 cycles of 95° C for 15 sec, 46° for 30 sec, and 70° C for 60 sec, followed by a final elongation of 70° C for 7 minutes. PCR products from the initial PCRs were diluted 25:1 in sterile H2O and we used this diluted product as template for an indexing PCR to add unique dual indexes to each sample for multiplexing. Unique Dual Indexing (UDI) primers (Integrative DNA Technologies, Coralville, Iowa, USA) contained the complimentary Illumina flow-cell adapter sequence and an 8 bp unique dual index on each end (forward and reverse) for unique sample identification from multiplexed samples. Indexing PCRs reactions included 10 μl of Amplitaq Gold DNA polymerase (Thermo Fisher Scientific), 0.5 uM of forward and reverse UDI indexing primers, and 2 μl of diluted PCR product from the first PCR in a total reaction volume of 20 μl. PCR cycling conditions for the indexing PCR were: initial denaturation at 95° C for 10 min, 15 cycles of 95° C for 15 sec, 50° C for 30 sec, and 72° C for 30 sec, followed by a final elongation of 72° C for 7 minutes. Following the indexing PCR, we cleaned up and normalized PCR products using a SequalPrep™ normalization plate (Thermo Fisher Scientific) following the manufacturer’s protocol. We then pooled the resulting product and provided it to the Center for Integrative Biosystems sequencing core at Utah State University where it was sequenced on an Illumina NextSeq 2000 with a 600 cycle P1 reagent kit.

Bioinformatic processing of MiSeq sequence data proceeded as follows. After demultiplexing, we removed primer sequences with CUTADAPT v. 1.18 (Martin 2011). Next, data were filtered, denoised, paired-ends were merged, chimeras were removed with DADA2 (Callahan *et al*. 2016), and Amplicon Sequence Variants (ASV) were merged into Operational Taxonomic Units (OTUs) at 97% similarity with VSEARCH (Rognes *et al*. 2016), all within the QIIME2 environment (Bolyen *et al*. 2019) version 2024.10. For taxonomic assignment of OTUs, we used the Python package BOLDigger3 (Buchner and Leese 2020), version 2.1.2, a classifier built specifically for taxonomic assignment of COI sequences using the Barcode of Life Database (BOLD). We ran BOLDigger3 with default parameters. We also performed taxonomic assignment with BLAST (Ye *et al*. 2012), and consensus between BOLDigger3 and BLAST was used for ultimate species assignment (Table 2). Assignment was made at species level if either BOLDigger3 or BLAST had a match percentage over 98%. If confidence was lower than 98%, we used the more conservative BOLDigger3 assignment to genus or family.

**Table 2.**
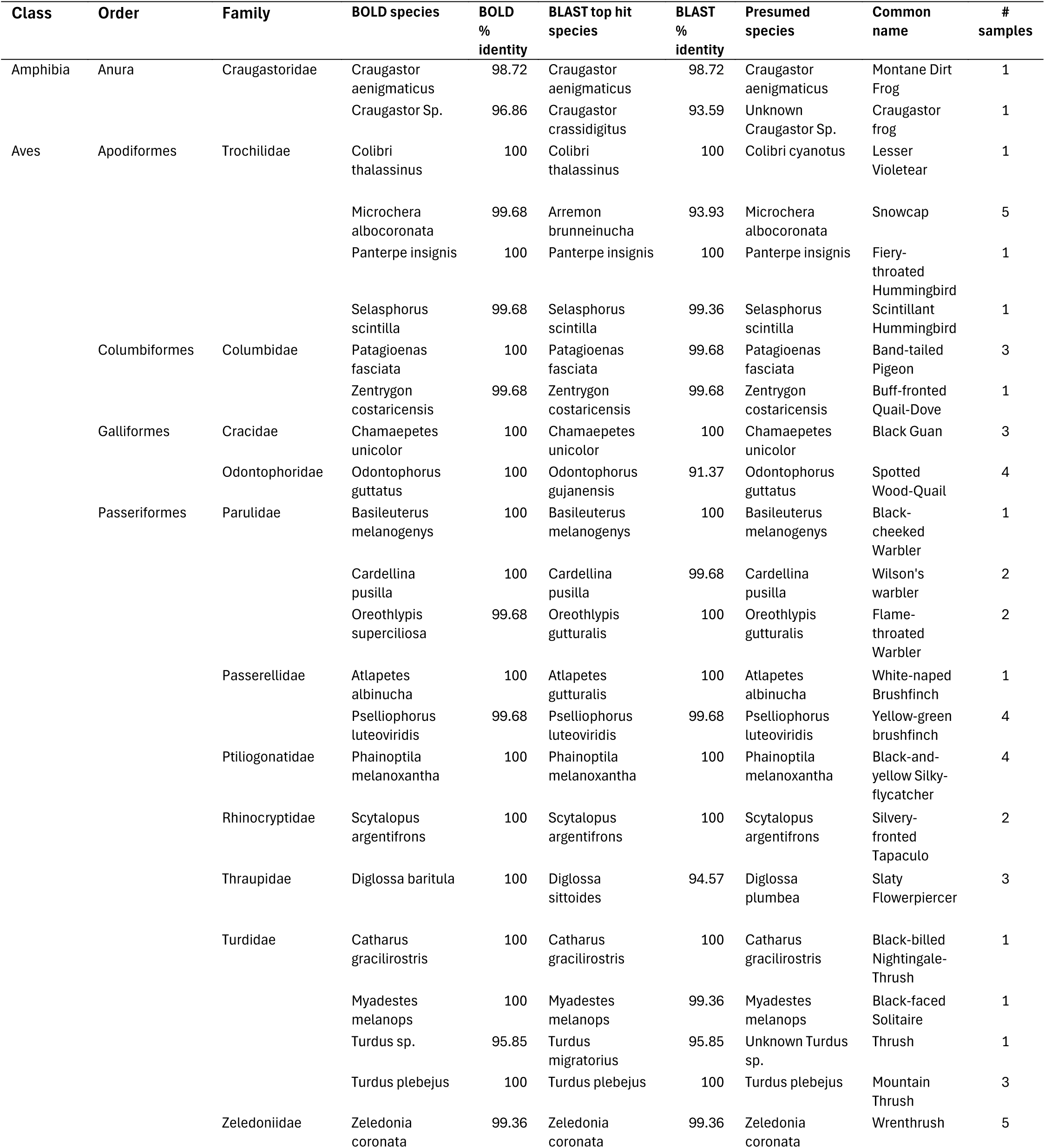
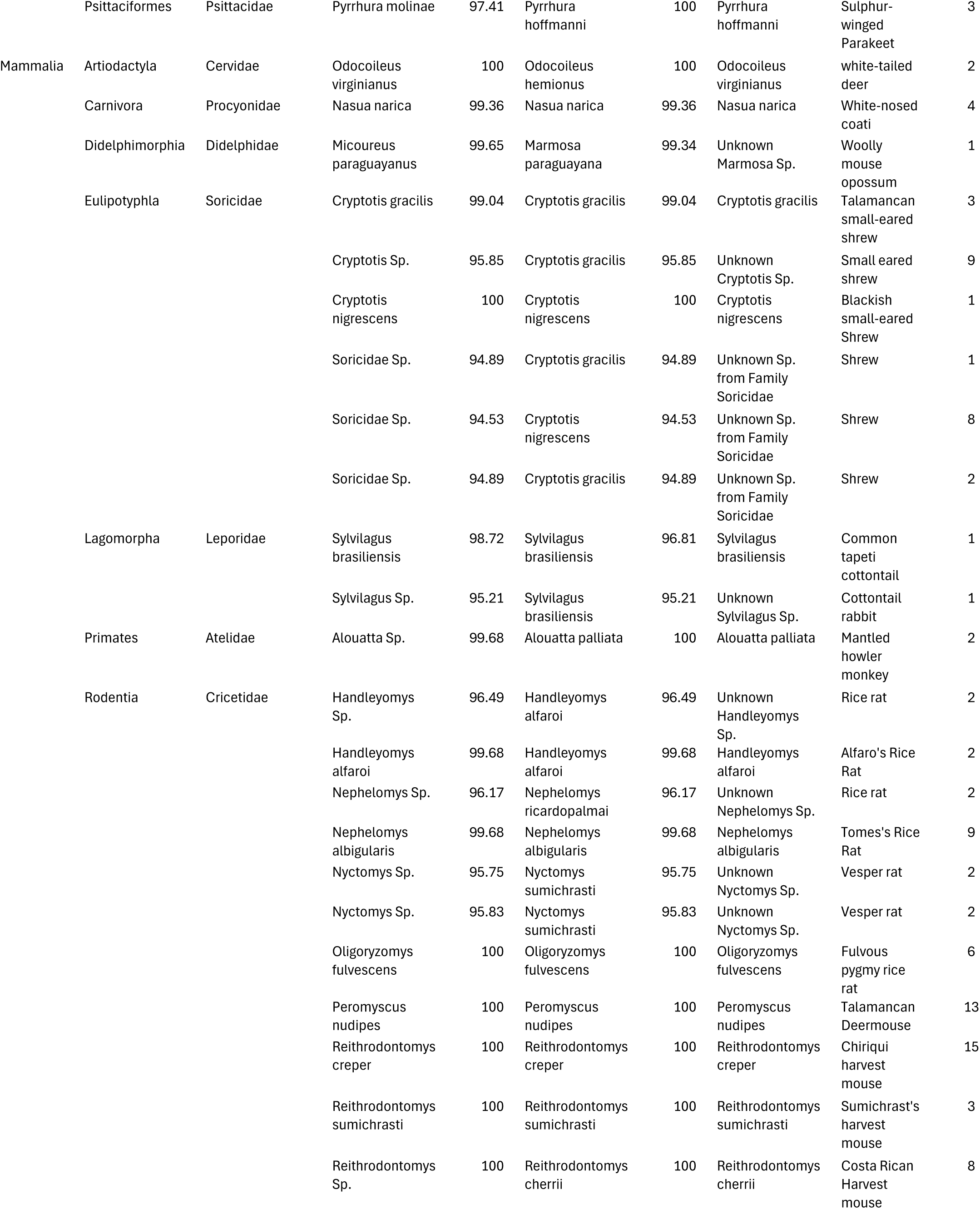
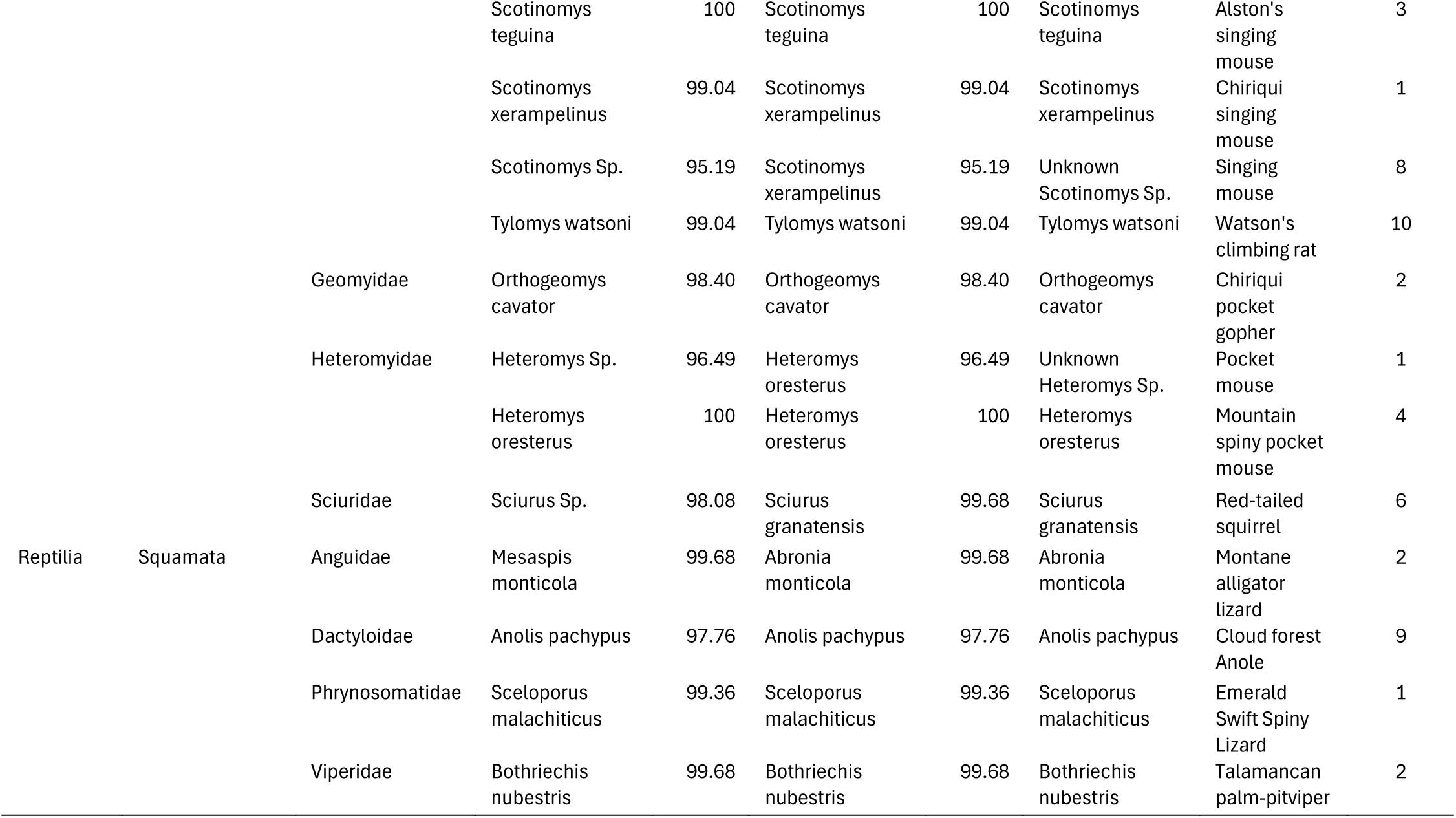
Prey species detected from DNA metabarcoding of *Leopardus pardinoides oncilla* scat samples collected in Panama and Costa Rica.

To estimate body mass distribution of prey species, for mammals and birds we used adult body mass estimates from EltonTraits 1.0 (Wilman *et al*. 2014). For species not present in the EltonTraits 1.0 dataset, body mass estimates were derived from median adult mass of museum specimen records accessed via VertNet (Constable *et al*. 2010) and GBIF (Global Biodiversity Information Facility, 2025; Supplementary table S2). For taxa only assigned to genus, we used median body mass of all members of the genus from EltonTraits 1.0 (Wilman *et al*. 2014). Because the reptiles and amphibians we detected are rare endemics, body mass estimates for these species were not available. Thus, we estimated body mass from length-mass relationships from published snout-vent length for these species (Arias et al., 2018; Doan et al., 2016; Meiri, 2008; Poe & Ibáñez, 2007; Supplementary table S2). Possible but improbable large prey species (*Odocoileus virginianus, Nasua narica, Alouatta palliata*; see discussion) were not included in prey body mass estimates.

## Results

### Distribution

We collected 421 potential *L. p. oncilla* scat samples from the field for analysis. Of these, 195 were confirmed as belonging to *L. pardinoides* by Sanger sequencing, 126 from Costa Rica, and 69 from Panama (Figure 1, Table 1). The remaining samples either did not amplify in PCR, likely due to DNA degradation, or were identified as non-target species such as Puma (*Puma concolor)*, Grey Fox (*Urocyon cinereoargenteus),* or Margay (*Leopardus wiedii).* We collected 107 (55%) confirmed *L. p. oncilla* samples by visual sampling, and 88 (45%) with the detection dog. We collected confirmed *L. p. oncilla* samples from Chriquí, Bocas del Toro, and Veraguas provinces in Panama; and Limón, Puntarenas, San José, Cartago, Heredia, and Alajuela provinces in Costa Rica (Table 1, Figure 1). We collected most *L. p. oncilla* samples (55%) from La Amistad International Park and adjacent private reserves. We collected 18% of confirmed *L. p. oncilla* samples from Volcán Barú National Park in Panama, 13% from the Cerro de la Muerte region of Costa Rica, including Tapantí-Cerro de la Muerte Massif National Park, Los Quetzales National Park, and several private reserves in the region, and 10% from Chirripó National Park. We collected 5 confirmed *L. p. oncilla* samples (2.6%) from the Central Volcanic Cordillera of Costa Rica including Turrialba National Park, Braulio Carrillo National Park, and Juan Castro Blanco National Park, and a single sample was collected from Santa Fe National Park in Veraguas Province, Panama (Table 1, Figure 1).

**Figure 1.**
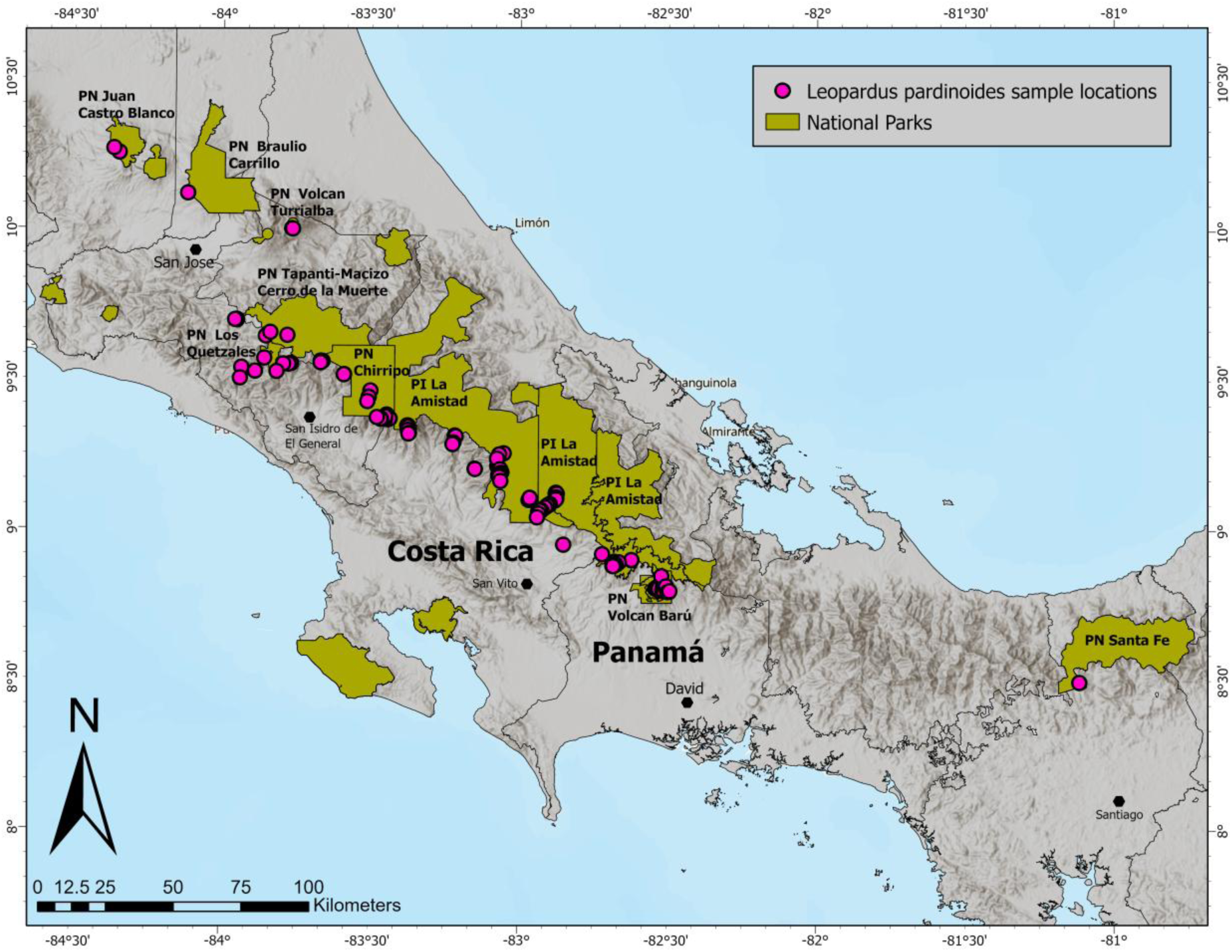
Locations of genetically confirmed *Leopardus pardinoides oncilla* scat samples collected from Panama and Costa Rica.

Median elevation of confirmed *L. p. oncilla* samples was 2805 m (range 821-3422 m). Just two confirmed samples were found from below 1700 m (1%), with 19 samples (10%) from below 2000 m, 117 (60%) from between 2000-3000 m, and 59 (30%) from above 3000 m (Figure 2). No confirmed *L. p. oncilla* samples were collected from the Tilarán mountains, or the Guanacaste Volcanic Cordillera in Costa Rica. Likewise, no samples collected from the Cerro Chucantí private nature reserve in the Majé mountains of Darien province, Panama were confirmed as belonging to *L. pardinoides* (Table 1).

**Figure 2.**
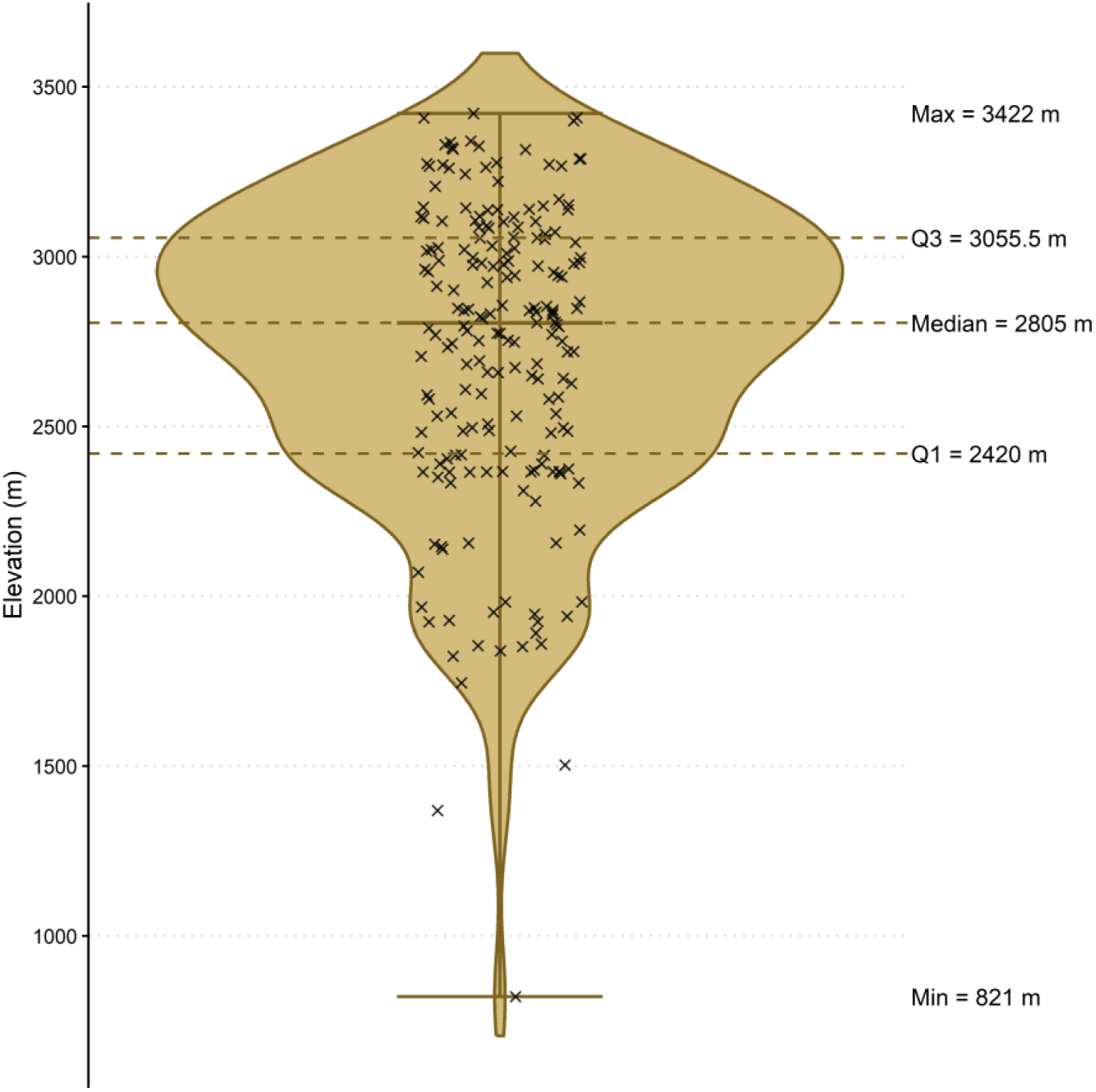
Distribution of elevation values for genetically confirmed *Leopardus pardinoides oncilla* scat samples (n = 195). X’s represent individual scat detection locations. The shaded violin represents kernel density estimate of elevations.

### Diet Metabarcoding

NextSeq sequencing of scat samples resulted in 102,769,738 reads, of which 23,089,742 reads were retained after quality filtering, denoising, and chimera removal, with a median of 113,526 reads per sample. Of these reads, 8,101,827 (35%) were assigned to *L. pardinoides*, and 604,345 reads (2.6%) were assigned to vertebrate prey taxa. The remaining reads were assigned to a variety of non-target taxa including fungi, bacteria, and arthropods, primarily from the orders Diptera and Coleoptera (Supplementary Table S3).

Of the 190 confirmed *L. pardinoides* samples used for diet metabarcoding, 106 samples (56%) contained vertebrate prey DNA sequences cumulatively belonging to 59 taxa: 31 mammals, 22 birds, four reptiles, and two amphibians (Table 2). Of the 106 samples containing prey sequences, 78 samples (74%) contained prey sequences of mammals, 37 (35%) of birds, 14 (13%) of reptiles, and two (1.9%) of amphibians (Figure 3). Among mammals, we found sequences belonging to seven orders and ten families, with rodents from the family Cricetidae (New Worlds rats and mice) the most common (55% of total samples), followed by species from the shrew family Soricidae (22% of total samples). Among birds, we found sequences belonging to five orders and 12 families, the most common being order Passeriformes (17% of total samples). The most common family was hummingbirds from the family Trochilidae (order Apodiformes, 7.5% of total samples), followed by Turdidae (5.7 % of total samples; Figure 4). For reptiles, all four taxa detected were from the order Squamata, comprising four families including Viperidae and three lizard families (Table 3). For Amphibia, both taxa detected were Anurans from frog genus *Craugastor*.

**Figure 3.**
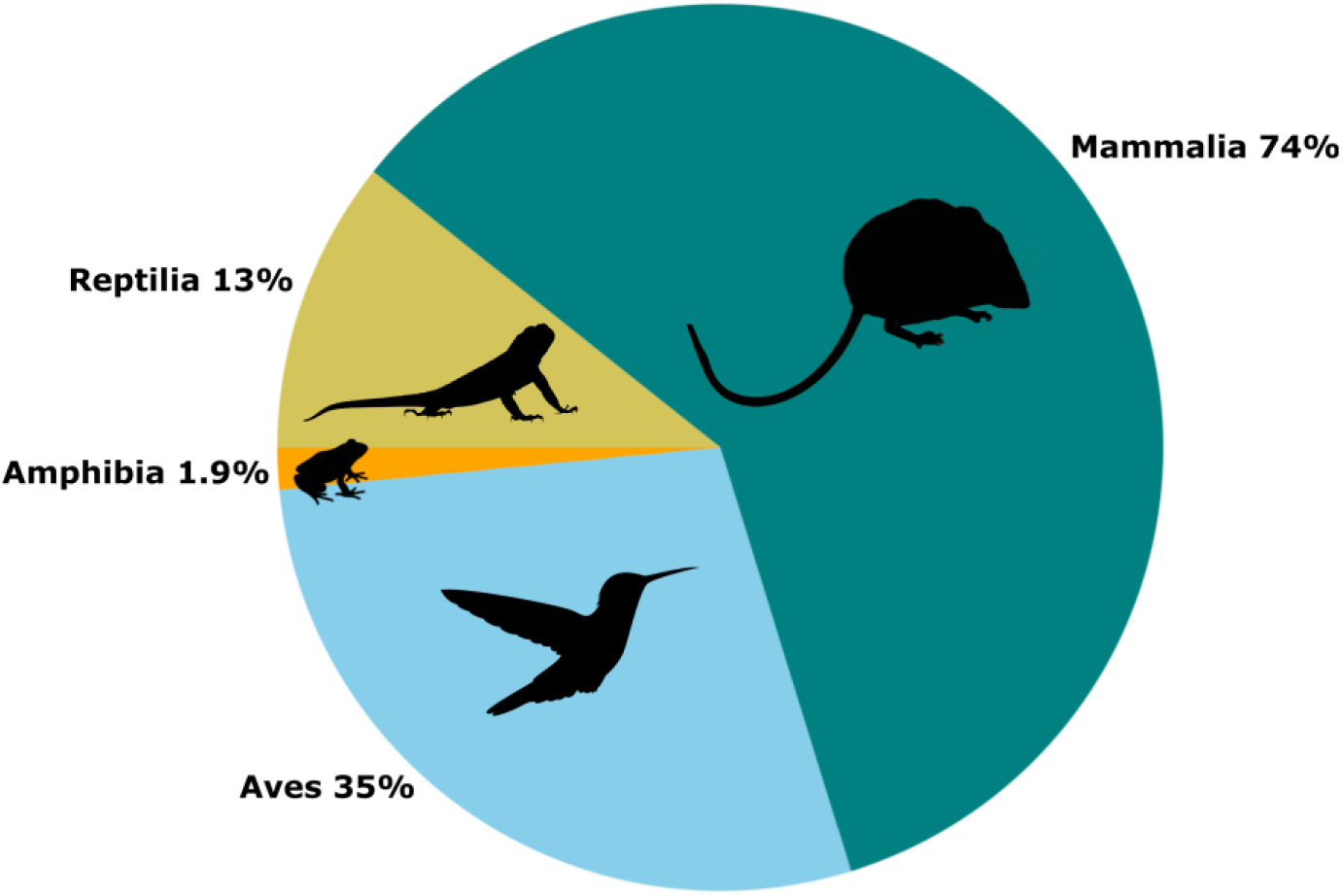
Proportion of scat samples containing prey DNA from each vertebrate class from metabarcoding of *Leopardus pardinoides oncilla* scats from Panama and Costa Rica.

**Figure 4.**
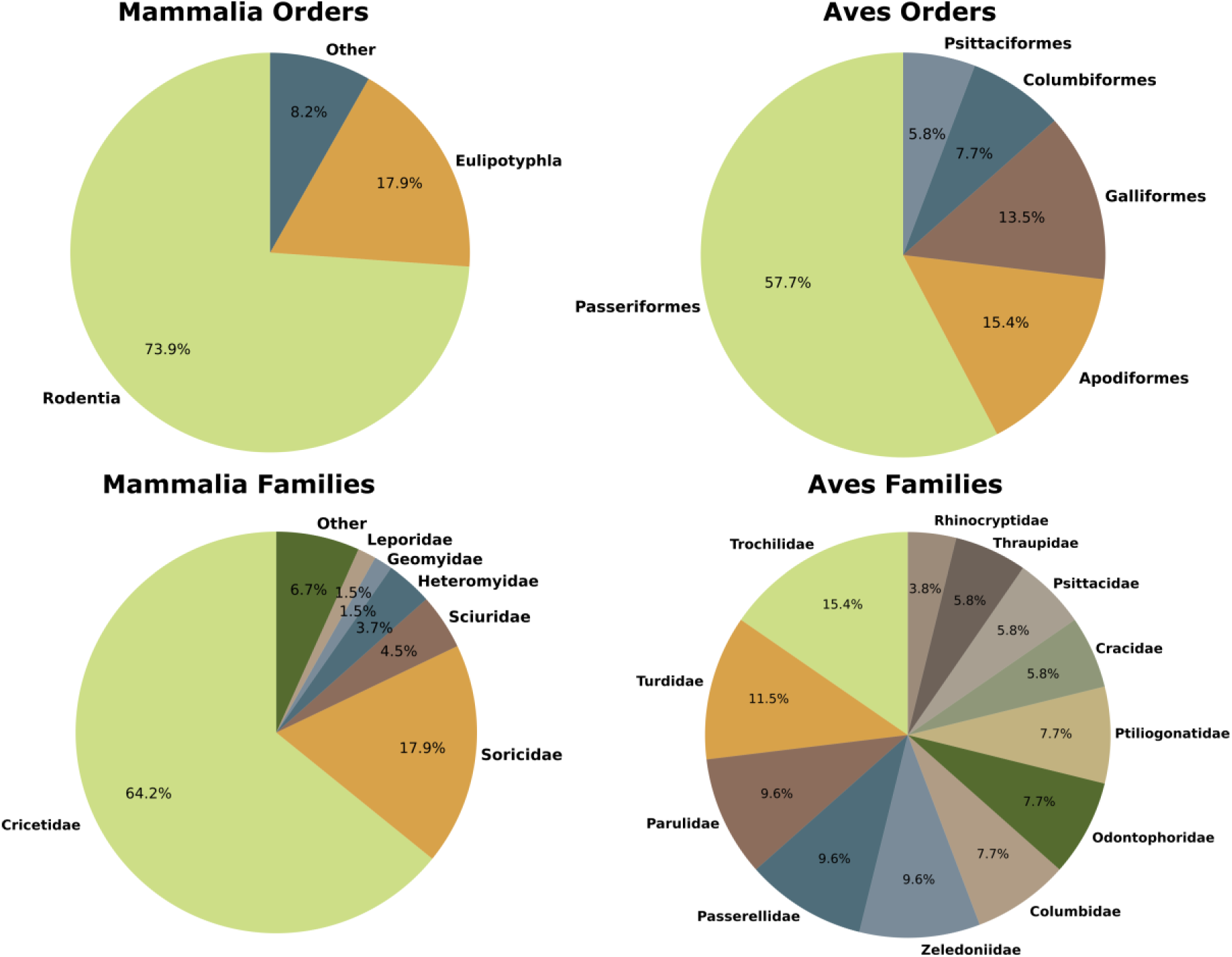
Proportional composition of *Leopardus pardinoides oncilla* diet from DNA metabarcoding. Pie charts show percentage of scats with prey DNA from orders and families of Mammalia and Aves.

Of the 59 taxa detected from *L. p. oncilla* scats, we were able to identify 76% to species level, 19% to genus level, and in 3 cases (5%) only to family level (Table 2). All three family-level assignments were taxa from the Family Soricidae. For birds, 95% of taxa were assigned to species level, and just one to genus level. A bird species that we assigned at species level, the Cinnamon-bellied Flowerpiercer *Diglossa baritula,* is not known from Costa Rica or Panama. However, the congener *Diglossa plumbea*, the Slaty Flowerpiercer, is known from our study area, but does not possess a reference sequence in BOLD. Likewise, the Mexican Voiletear *Colibri thalassinus* is not known from Costa Rica or Panama, but the Lesser Violetear *Colibri cyanotus* occurs in our study area but does not possess a publicly available reference sequence. Thus, we believe the sequences we observed from these genera likely came from *D. plumbea* and *C. cyanotus.* For mammals, 61% were assigned to species level, 29% to genus level, and 10% to family level. For reptiles, all four taxa were assigned to species level, and for amphibians, one was assigned to species, and one to genus.

Estimated median adult body mass for detected prey species was 25.0 g (range 1.5-1343 g). For mammals, median adult body mass was 46.2 g (range 7-1343 g), for birds 25.5 g (range 2.2-1135 g), for reptiles 16.3 g (range 3-230 g), and for amphibians 1.5 g (range 1.5-1.5 g; Figure 5, Supplementary Table S2).

**Figure 5.**
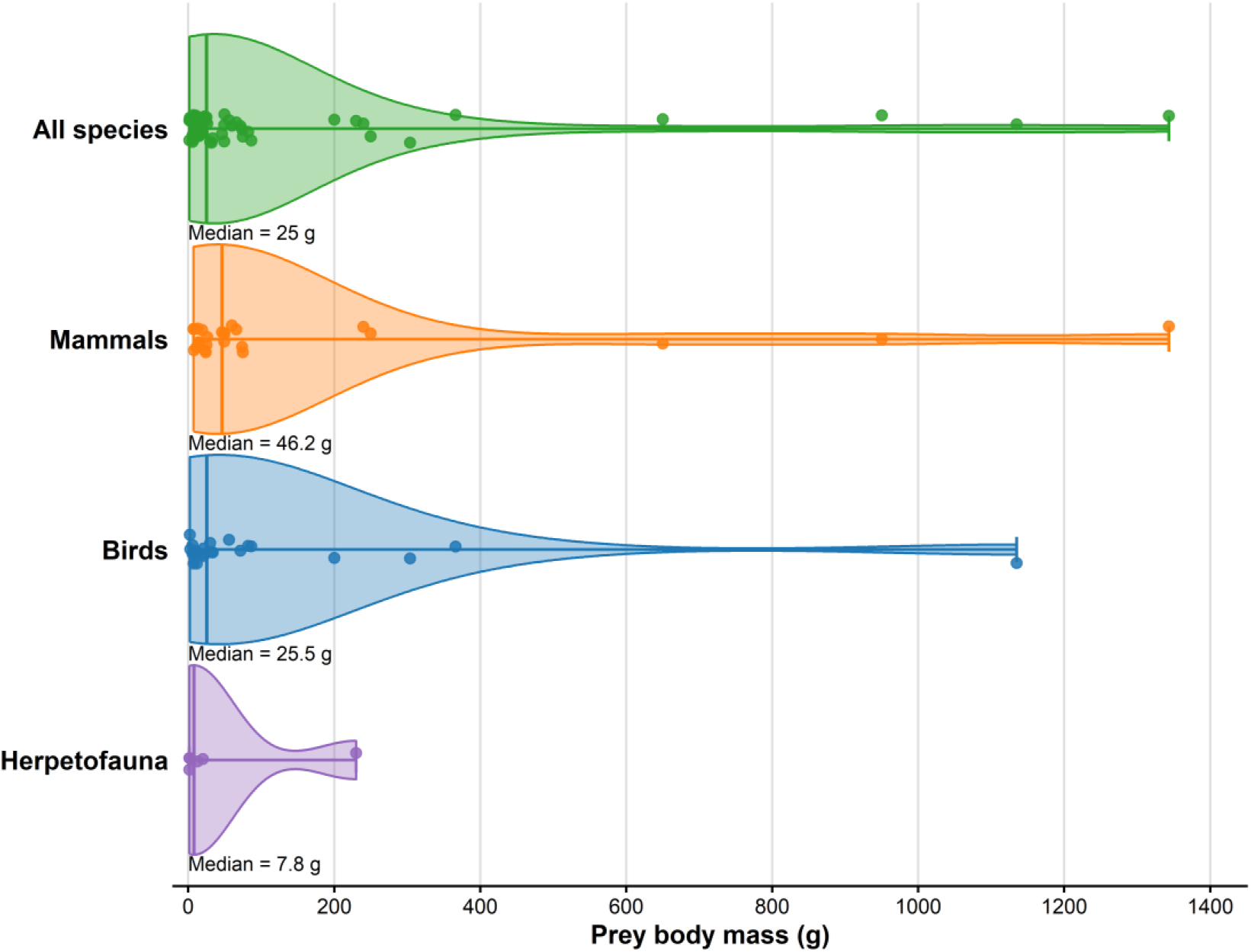
Distribution of estimated adult body mass of prey detected from DNA metabarcoding. Shaded areas show kernel density estimates, points represent individual prey taxa, and vertical lines indicate median values.

## Discussion

This work makes a substantial contribution to our understanding of the distribution of the Clouded Tiger Cat *L. pardinoides oncilla* in Central America, and we provide the first comprehensive evaluation of the diet of *L. pardinoides*. Information on the ecology and natural history of this species is both important and timely since *L. pardinoides* was recently recognized as a new species that faces imminent threats from habitat loss (de Oliveira et al., 2024). Our results also strongly support the restriction of *L. p. oncilla* to high elevation montane habitat in Central America. This suggests that this species and its prey base is likely to face future impacts from climate change due to disproportionate effects on montane predator and prey communities from elevational dependent warming (Pepin *et al*. 2015) and upslope range compression (Freeman *et al*. 2018).

### Distribution

Previous studies relying on camera trapping have contributed to our knowledge of the distribution of Clouded Tiger Cats throughout their range, including Central America (de Oliveira et al., 2024; Ramírez-Fernández et al., 2024). However, our results substantially expand the known distribution of Clouded Tiger Cats in Central America by more than doubling the number of known records from Central America while providing records from new locations. Many of the new records are from Panama, where the species has been less studied than in Costa Rica.

All our verified records exist within the model-predicted species distribution of de Oliveira et al. (2024). However, confirming empirical species presence within this predicted range is important for conservation efforts. The vast majority (97%) of our *L. p. oncilla* records come from the Talamanca Cordillera mountains in Costa Rica and Panama and from the Cerro de la Muerte massif in Costa Rica, ranging from Volcan Baru National Park in the north to Panama in the south, and including protected areas in between (Table 1, Figure 1). This region, stretching for approximately 155 km across the international border along the continental divide of Central America, includes a series of protected areas containing optimal high elevation *L. p. oncilla* habitat. These protected areas include Los Quetzales National Park, Tapantí - Cerro de la Muerte Massif National Park, Chirripó National Park, La Amistad International Park, Volcan Baru National Park, and several connected private reserves which are contiguous, all providing a large area that is the core of Clouded Tiger Cat habitat in Central America.

Due to accessibility, most of our field work, and thus most of our records, are from the Pacific side of the continental divide. However, large areas of suitable habitat exist on the Caribbean side of the divide, particularly in La Amistad International Park, an area less impacted by anthropogenic disturbance due to its remoteness. Although our records are presence only, and not species abundance or density, the highest densities of *L. p. oncilla* scat samples were collected from the highest and most remote regions of La Amistad International Park, including the Cerro Pittier and Cerro Fabrega area along the Costa Rica-Panama border where we collected 38 samples in just 5 days of visual sampling, and the Cerro Kamuk area of Costa Rica where we collected 27 samples in five days of sampling with a detection dog. Thus, we believe these areas, and adjacent locations on the Caribbean side of the continental divide, likely contain some of the highest densities of the species in Central America.

We also collected a relatively large number of samples, albeit with considerably more effort, from Chirripo National Park in Costa Rica (n=20 samples in 13 days), and Volcan Baru National Park in Panama (n=36 samples in 20 days; Table 1). These parks experience considerably more human presence from tourism and recreation because they contain the highest points in their respective countries. Still, pressures from tourism and recreation only impact a portion of each park, so these activities may have minimal impact on *L. p. oncilla* populations. Finally, we collected a relatively large number of samples from the Cerro de la Muerte region of Costa Rica including Los Quetzales National Park, Tapantí - Cerro de la Muerte Massif National Park, and adjacent private reserves, albeit with substantial effort mostly with a detection dog (n=25 samples in 28 days; Table 1). This region sees even more anthropogenic impact, not only from tourism and recreation, but because it is traversed by Costa Rica’s Interamerican Highway, one of the busiest highways in Costa Rica (González-Maya *et al*. 2024), with 3,400 vehicles per day reported in 2021. It is unclear how much impact the highway has on Clouded Tiger Cat movement, although numerous *L. p. oncilla* roadkills have been recorded here. Efforts led by the NGO Panthera are ongoing in this area to modify culverts as temporary wildlife crossing tunnels to aid animal movement under the highway until permanent wildlife crossing can be built.

We collected just five *L. p. oncilla* samples (2.6%) from outside of the Talamanca mountains, all but one with the scat detection dog (Table 1). These samples were collected from the Central Volcanic Cordillera of Costa Rica, from three different national parks, Turrialba, Braulio Carrillo, and Juan Castro Blanco. Although we were able to confirm the presence of the species in these parks, it is likely that Clouded Tiger Cats face the greatest conservation threats in the Central Volcanic Cordillera due to habitat loss. The parks where we detected *L. p. oncilla* are relatively small, isolated from one another, and very close to substantial population centers (Pedroni *et al*. 2008). Because considerable *L. p. oncilla* habitat in this region has been lost to cattle ranches, dairy farms, roads, and other forms of agriculture, including some habitat inside of, and connecting protected areas, populations in this region may face the most immediate threats of local extinction, and thus may warrant the greatest conservation efforts and research to estimate population sizes, threats, and viability.

Finally, we collected a single *L. p. oncilla* sample from Santa Fe National Park in the Cordillera Central of Panama, the far eastern end of the greater Cordillera Talamanca, approximately 150 km to the east of most known records. Although a single previous camera trap record existed from this region at an unusually low elevation at the edge of the National Park (de Oliveira et al., 2024; Fox-Rosales, personal communication), this study, along with another recent camera trap study (Rodgers *et al*. 2026), confirms the presence of Clouded Tiger Cat in Santa Fe National Park. Our scat detection was from near the summit of a mountain at 1369 M, one of the lowest elevation samples we collected. Given this finding, it is likely that *L. p. oncilla* also occurs in the Central Cordillera between Sante Fe National Park and the nearest known records to the west in Volcan Baru National Park approximately 150 km away. This area falls mostly within the Comarca Ngäbe-Buglé, an autonomous Indigenous administrative region that possesses habitat up to 2500 m in elevation, with no confirmed records of *L. p. oncilla* albeit with no past systematic surveys.

There were also several areas where we failed to detect the presence of *L. p. oncilla*. The first was the Tilarán mountains, where we conducted visual searches within three private reserves: Monteverde Cloud Forest Reserve, Santa Elena Cloud Forest Reserve, and the Children’s Eternal Rainforest in Puntarenas province of Costa Rica. Many verified records of Clouded Tiger Cats have been obtained from this region, and the species certainly occurs here despite our failure to find samples (Rogan 2021; Ramírez-Fernández *et al*. 2024). As this area falls within the upper elevation limit for ocelots, it is possible that samples were more difficult to find because Clouded Tiger Cats change their marking behavior along trails to be less conspicuous (de Oliveira et al. 2009, 2022), making visual searches for scat samples difficult. We also failed to find any samples in the Guanacaste Volcanic Cordillera in Costa Rica, where we conducted visual searches in Miravalles National Park and Rincón de la Vieja National Park. These protected areas occur in relatively high elevation cloud forest habitats that seem suitable for *L. p. oncilla* (up to 2000 m in Miravalles, and 1900 m in the Volcan Santa Maria zone of Rincón de la Vieja National Park), and they lie within the envelope of predicted suitable habitat from modeling by de Oliveira et al. (2024). However, these parks are isolated and do not have a high elevation connection with other known *L. p. oncilla* populations. Our searches were by no means exhaustive, so although it is possible that *L. p. oncilla* does not occur in Guanacaste province due to isolation or competition, more comprehensive sampling with a detection dog or camera traps is needed to determine if the species is absent from this region.

We also did not find any *L. p. oncilla* samples from the Majé Mountains of Darien Province in Panama, within Chucantí Private Nature Reserve. We collected four small putative felid fecal samples here, but two did not yield DNA, and the other two were identified as Margay. This location has a small area of cloud forest habitat which is extremely isolated, with a maximum elevation of 1439, yet it was identified by de Oliveira et al. (2024) as potentially suitable for Clouded Tiger Cats. This location is approximately 300 km distant from the nearest known *L. p. oncilla* records from Panama, and a slightly greater distance from the nearest records in Colombia (de Oliveira et al., 2024). The Darien region of Panama has been recognized as a potential barrier to historical dispersal of Clouded Tiger Cats from Colombia to Central America due to its low elevation (Bonilla-Sánchez et al., 2024), leading to genetic differentiation between South American and Central American populations (Lescroart *et al*. 2023). Méndez-Carvajal & Gutiérrez-Pineda, (2024) report detection of *L. p. oncilla* from Chucantí Private Nature Reserve with camera trapping. We have not reviewed all their alleged *L. p. oncilla* photographs as they are not publicly available, however, we believe the single photograph accompanying the publication is not sufficient proof that *L. p. oncilla* occurs here. This photograph could certainly be of *L. p. oncilla*, however given the difficulty of distinguishing *L. p. oncilla* from *L. wiedii or* juvenile *L. pardalis* (which also occur here) from photographs, we believe that greater evidence is needed to conclusively say that Clouded Tiger Cats occur in Chucantí, such as genetic evidence, or photographs of a melanistic individual. Melanism is not known for *L. wiedii* or *L. pardalis*, but it occurs in up to 32% of *L. p. oncilla* individuals in populations from Costa Rica (Mooring *et al*. 2020). We are also not aware of any verified *L. p. oncilla* records that occur at such low elevation when not connected to higher elevation areas of at least 1800 m.

Consistent with previous research (de Oliveira et al., 2024; Ramírez-Fernández et al., 2024), we found that *L. p. oncilla was* strongly associated with high elevation cloud forest habitats, mostly above 1700 m. The median elevation for our confirmed Clouded Tiger Cat samples was 2805 m, with greater than 75% of samples found above 2400 m (Figure 2). Samples were found near some of the highest points in Costa Rica and Panama, including a sample from 3422 m in Cerro Chirripó, the highest mountain in Costa Rica, from 3409 m just below the summit of Volcan Baru, the highest mountain in Panama, and from 3341 m just below Cerro Fabrega, the second highest mountain in Panama. Just two samples were found below 1700 m, the first the aforementioned sample from Sante Fe National Park. We also found a single sample from just 821 m in an area southwest of Cerro de la Muerte, confirming that the species can be found at elevations below 1000 m. However, since this sample is a significant outlier, it seems plausible that it was from a dispersing individual searching for suitable habitat. The sample was found in a forested patch with open pasture lands on either side, however, with forested connection to higher elevation habitat, with elevations of 1500 m just 2.6 km away, and elevations of 2000 m 3.6 km away.

Our finding of most samples at high elevations may have been somewhat influenced by confirmation bias. Visual searches often targeted high elevation areas that we believed would be suitable habitat for Clouded Tiger Cats based on existing data. However, in most cases, to get to these high elevation habitats, we began walking well below 2000 m (Table 1). Thus, we searched lower elevations on the way to the higher elevation areas where most of our samples were found. In addition, the scat detection dog team conducted 81 days of surveys from low elevation areas below 1000 m, primarily searching for Jaguar *Panthera onca* samples, but no *L. p. oncilla* samples were found during these lower elevation searches (Supplementary Table S1). Thus, confirmation bias likely had minimal influence on the elevations where samples were found. It is possible that our high elevation findings reflect habitat availability. Although there is much less area above 2000 m in Costa Rica and Panama than below 2000 m, much of the area between 1000-2000 m is prime agricultural land that has been converted to cattle ranches, dairy farms, and cash crops such as coffee. Above 2000 m, terrain is often more rugged, and less suitable for agriculture. Thus, it is possible that Clouded Tiger Cats were more abundant below 2000 meters historically but have been forced to higher elevations due to habitat loss at lower elevations. Below 1500 m, competition with Ocelots may also be a limiting factor independent of habitat suitability (de Oliveira *et al*. 2009, 2022). We conducted very little sampling in agricultural landscapes; however, we did collect one *L. p. oncilla* sample from a cattle pasture at the edge of forest in La Amistad International Park in Panama (some agricultural and cattle ranching is still permitted on land converted to agriculture prior to the creation of National Parks in Panama and Costa Rica), and we also found one sample in a forest less than 25 meters from a coffee field in a private reserve bordering La Amistad International Park, and another just 30 m from and agricultural field in Volcan Baru National Park. Future research should investigate Clouded Tiger Cat use of agriculturally modified landscapes.

We recorded numerous records of *L. p. oncilla* in the open páramo ecosystem above tree line near the tops of mountains where camera trap studies are rare because of the difficulty of deploying cameras in treeless, high elevation environments. Although Gonzalez-Maya and Schipper (2008) recorded *L. p. oncilla* from the cloud forest - páramo transition zone in Chirripó National Park with camera trapping, most past research on *L. p. oncilla* describes the sub-species as an upper montane or cloud forest specialist (Ramírez-Fernández et al., 2024). Indeed, the vast majority of our records are from cloud forest habitats, but we did encounter 11 *L. p. oncilla* samples in páramo on Cerro Fabrega, the largest area of páramo in Panama within la Amistad International Park, as well as in Chirripo National Park, the highest mountain and largest páramo in Costa Rica, and in Volcan Baru National Park, the highest mountain in Panama. This does not verify that páramo habitat is important to *L. p. oncilla*; it is possible that animals are simply crossing páramo to move between patches of cloud forest on opposite sides of the continental divide. However, it does provide evidence that páramo is not a barrier to movement or dispersal for *L. p. oncilla* and might be significant habitat for the species

### Diet Metabarcoding

We found that *L. p. oncilla* has a generalist diet; 59 different prey taxa were detected from metabarcoding of scat samples. Similar to *L. tigrinus* diet studies (Silva-Pereira et al., 2011; Trigo et al., 2013; Wang, 2002), we found that the *L. p. oncilla* diet was dominated by small mammals, with a substantial part of the diet also consisting of avian prey. Reptiles were also frequent, but less common, and anurans were infrequently detected (Table 2, Figure 3). Amongst mammal diet items, cricetid rodents were most common, followed by shrews, and then rodents from the orders Heteromyidae and Sciuridae (Figure 4). Amongst birds, songbirds from the order Passeriformes were the most common, but hummingbirds were also frequently detected (Figure 4). Bird prey was from a wide range of families (12 in total). Amongst Reptilia, lizards were detected from 12 samples, most frequently the endemic and recently described cloud forest anole lizard *Anolis pachypus* (Poe and Ibáñez, 2007). Astonishingly, we also detected Talamancan Palm-pit viper *Bothriechis nubestris* in the diet, a recently described venomous snake species endemic to the cloud forests of the Cordillera Talamanca (Doan et al., 2016). From two samples, we detected two frog taxa, both from the genus *Craugastor,* including the recently described endemic species *Craugastor aenigmaticus* (Arias et al., 2018).

We detected sequences from three mammal species that we believe may or may not be prey items for *L. p. oncilla*. We detected DNA in four samples from the common procyonid species White-nosed Coati *Nasua narica*. It is certainly plausible that a Clouded Tiger Cat preyed on an infant or juvenile Coati, but consumption of an adult Coati seems unlikely given that adult Coatis are larger in size than *L. p. oncilla*. *Nasua narica* has been known to roll in the feces of other carnivores, and to rub on Ocelot latrines (Rodgers *et al*. 2015; King *et al*. 2017; Fleming and Weldon 2021). Likewise, we found sequences from Mantled Howler Monkey *Alouatta palliata* from two samples. Once again, predation of a baby Howler Monkey by a Clouded Tiger Cat is not impossible, but seems unlikely, as Howler Monkey mothers are highly vigilant in protecting infants (Treves *et al*. 2003). Howler Monkeys are common in the area where we detected their DNA, and they frequently urinate and defecate from the canopy above, so it is possible that DNA from above landed on our samples. For both Coati and Howler Monkey, reads counts were very low in all samples, and thus natural contamination of samples in the field with Coati or Howler Monkey DNA seems more likely than predation by *L. p. oncilla*. We also detected sequences from White-tailed Deer *Odocoileus virginianus* from two samples, including one with relatively high read counts from Volcan Barú National Park. *Odocoileus virginianus* occurs in Volcan Barú National Park, but due to the size difference it would be impossible for a Clouded Tiger Cat to hunt even an infant White-tailed Deer. However, it is plausible that the *O. virginianus* sequences we obtained were from *L. p. oncilla* scavenging a carcass. Facultative scavenging is not uncommon among felid species, and scavenging on deer carcasses has been well documented for the small cat species *Felis silvestris* (Krofel *et al*. 2021).

The lower percentage of prey taxa assigned to species level for mammals (61%), vs. birds (95%) reflects the reality that reference sequence databases for small mammals, especially shrews, is incomplete from remote, upper elevation cloud forest regions of Central America. Neotropical shrew species, particularly from the genus *Cryptotis*, are particularly difficult to identify based on morphology. Cryptic species of *Cryptosis* likely exist that have not yet been adequately described (Guevara *et al*. 2024), and new species have been recently discovered (Guevara *et al*. 2014; Guevara 2023) including from Costa Rica (Woodman and Timm 2017). Thus, it is possible that some of the five unidentified shrew species in the diet of *L. p. oncilla* are undescribed cryptic endemics.

For taxa that we were able to assign to the species level, the majority (58%) are endemic to the mountains of Panama and Costa Rica. This included 42% of detected mammals, 67% of birds, 75% of reptiles, and 100% of amphibians. It is likely that many of the taxa we could only assign to genus or family are also endemic species that lack reference sequences. High elevation Neotropical cloud forests are hot spots for endemism (Almendra and Rogers 2012; Jankowski *et al*. 2021; Ramírez-Fernández *et al*. 2024). Since most of the species eaten by Clouded Tiger Cats live nowhere else on earth, conservation of Clouded Tiger Cats in Central America will require conservation of the many endemic species that make up their prey base.

We used the Leray–Geller COI primer pair mlCOIintF/jgHCO2198 (Leray et al. 2013; Geller et al. 2013) for metabarcoding because it targets the Cytochrome c oxidase I gene (COI), the most commonly used gene for barcoding of animals that is the best represented in reference databases (Ratnasingham and Hebert 2007; Kress *et al*. 2015). This primer pair has been used for numerous diet studies focused on marine and freshwater taxa (Leray *et al*. 2015; Matley *et al*. 2018), as well as for environmental DNA metabarcoding (Stat *et al*. 2017; Hajibabaei *et al*. 2019) but it has not been frequently used for carnivore diet metabarcoding (but see Havmøller et al., 2021). In the Neotropics, and especially from hyper-diverse and understudied areas like high-elevation cloud forests, reference databases are incomplete, especially for non-charismatic animals such as small mammals. However, reference databases are much more complete for COI than for other genes that are commonly used in carnivore diet metabarcoding studies. The most commonly used marker for diet metabarcoding in carnivores targets the mitochondrial gene 12S, which is not nearly as well represented as COI (Riaz *et al*. 2011; Shehzad *et al*. 2012).

Additionally, we have found that although this 12S marker works well for mammalian prey, it has poor taxonomic discrimination power for birds and is often only able to assign bird taxa to order, especially within the order Passeriformes (Draper *et al*. 2022). As our goal was to detect a generalist diet of mammals, birds, reptiles, and amphibians, the COI marker we used was preferable, and given the wide breadth of taxa we detected from four different classes, we believe it performed well for this purpose.

The Leray–Geller COI primer pair we used (Geller et al., 2013; Leray et al., 2013) does have some minor drawbacks compared to other markers commonly used for diet analyses. First, as this primer set is highly degenerate, and designed to amplify nearly all metazoans rather than just vertebrates, it produces a large number of non-target sequences. Indeed, over 60% of our data were sequences from non-vertebrate species such as bacteria, fungi, and invertebrates. Of the invertebrate species detected (Supplementary Table S3), it is possible that some species were in fact eaten by *L. p. oncilla*. However, because of the high sensitivity of DNA metabarcoding, we believe that most of the invertebrate DNA amplified was environmental. Because it is impossible to determine if the invertebrate DNA we amplified was ingested by *L. p. oncilla*, or simply introduced from the environment, we will not speculate if invertebrates are an important part of the diet of Clouded Tiger Cats.

Because the Leray–Geller COI primer pair is highly degenerate, designing a predator host DNA blocking primer for use with this marker is not feasible. Many carnivore diet analyses use blocking primers to reduce amplification of DNA from the predator host so that more target prey DNA is amplified (Shehzad et al., 2012). Indeed, 35% of our data was read from *L. pardinoides*, and overall, just 2.6% of our data was from target vertebrate prey taxa. Fortunately, we anticipated this issue, and we conducted deep sequencing of our metabarcoding library. We sequenced the library on an Illumina NextSeq run that yielded approximately 100 million reads, instead of on the more common Illumina MiSeq platform (which has a maximum yield of 25 million reads). Due to this deep sequencing, we had enough vertebrate prey data (604,345 reads) to adequately describe the diet of Clouded Tiger Cats despite the majority of data being non-prey sequences.

### Conservation Implications

For the conservation of *L. p. oncilla*, core habitat in the Cordillera Talamanca must be vigilantly protected as a high elevation refugia for the sub-species. In the Central Volcanic Cordillera of Costa Rica, it is imperative to protect remaining high elevation habitat through the restoration of agricultural land back to cloud forest, especially in areas that provide connectivity between isolated volcanos. More detailed research on population abundance and viability in the Central Volcanic Cordillera is urgently needed to assess the risk of local extinction in isolated populations. Our results on the diet of *L.p. oncilla* demonstrate that they rely on many endemic prey species for survival. As such, Clouded Tiger Cats can act as an umbrella species for the conservation of endemic small mammals, birds, reptiles and amphibians that make up the amazingly diverse fauna of the montane cloud forests of Panama and Costa Rica.

## Acknowledgements

We thank Peter Robertson, Edgar Toribio Pérez, Deiver Espinoza-Muñoz, Elio Altamirano, Jonathan Morillo Chaves, Katherine Arauz-Ponce, Marlon Olmos, Floidany Ureña Jimenez, Amy Eppert, Steven Blankenship, Sierra Ullrich, and Abigail Wagner for assistance in the field. We thank Lindsay Stallcup from Bosque Eterno De los Niños, Yoryineth Mendez and Isabela Vargas from Monteverde Cloud Forest Reserve, and Natalia Vallegas from Santa Elana Reserve for help with logistics or field assistance. We thank SINAC/MINAE Costa Rica, and SINAC officials Roger Gonzalez Tenorio, Junior Porras, Enza Vargas, and Andres Mora. Finally, we thank the detection dogs Google and Tigre for their hard work in the field and Working Dogs for Conservation for providing detection dog handler training. Samples from Costa Rica were collected under permits R-CM-P-001-2023-OT-CONAGEBIO (ACLAP-ACG-ACHN-ACT-ALAC-ACOSA) and R-CM-P-002-2023-OT-CONAGEBIO (ACC-ACAT-ACOPA- ACTO). Samples from Panama were collected under Ministerio de Ambiente collection permits SE/A-4-20 and ARG-0144-2022.

## Author Contributions

TR designed the study, acquired funding, applied for Panamanian permits, conducted field and all laboratory work, and wrote the initial manuscript draft. RSP conducted field work, acquired funding, provided institutional and logistical support, applied for Costa Rican permits, and helped with manuscript revision and editing. SAA conducted field work and helped with manuscript revision and editing. EVA conducted field work. PLC conducted field work and helped with manuscript revision and editing. DAG provided logistical and institutional support, applied for Costa Rican permits, and helped with manuscript revision and editing. CMM conducted field work and helped with manuscript revision and editing. MM conducted field work and helped with manuscript revision and editing. MV helped with sample handling, applied for Panamanian permits, and provided logistical and institutional support. KM provided supervision, institutional support, and helped with manuscript revision and editing.

## Data Availability

All metabarcoding sequence data can be found on the NCBI Sequence Read Archive under BioProject PRJNA1331444.

## Supplementary Data

**Supplementary Table S1.** Locations sampled with a scat detection dog below 1000 m in elevation. Zero *Leopardus pardinoides* scat samples were encountered at these locations.

**Supplementary table S2**. Body mass estimates and sources for *Leopardus pardinoides* prey species detected from DNA metabarcoding of scat samples.

**Supplementary Table S3.** Arthropod species detected from DNA metabarcoding of *Leopardus pardinoides* scat samples.

