## Supplementary material for "Distribution and diet of Central American Clouded Tiger Cat *Leopardus pardinoides oncilla* via noninvasive genetics": supplimentary

**Supplementary Table S1.** Locations sampled with a scat detection dog below 1000 m in elevation. Zero *Leopardus pardinoides* scat samples were encountered at these locations.

| Location | Elevation Range (m) | # sampling days |
|---|---|---|
| Casas Negras, Toro Sentado, Finca Clemencio | 200 - 400 | 2 |
| Estación Biológica El Zota, RNVS Barra del Colorado | 0 - 50 | 2 |
| Estación Experimental Forestal Horizontes | 50 - 300 | 3 |
| Finca Chirripó | 500 - 1000 | 2 |
| Finca Eladio | 50 - 200 | 1 |
| Finca Maritza | 100 - 300 | 3 |
| Finca Río Banano | 0 - 50 | 2 |
| Finca Sandoval | 100 - 300 | 2 |
| Our Planet Foundation | 500 - 800 | 2 |
| Papagayo Peninsula | 0 - 50 | 21 |
| Parque Nacional Braulio Carrillo: El Ceibo | 100 - 600 | 3 |
| Parque Nacional Tortuguero | 0 - 120 | 20 |
| Playa Moin | 0 - 20 | 2 |
| Rancho Quemado, Península de Osa | 0 - 100 | 1 |
| Reserva Las Brisas | 150 - 250 | 1 |
| Reserva Natural Pacuare | 0 – 200 | 2 |
| Reserva Selva Bananito | 100 - 300 | 4 |
| Sendero El Tigre, Península de Osa | 0 - 100 | 1 |
| Senderos Ecológicos Finca San Gerardo | 800 - 1200 | 1 |
| Trocha Oreamuno-Guápiles | 50 - 150 | 1 |
| Universidad EARTH | 50 -100 | 2 |
| Veragua Rainforest | 50 - 200 | 3 |
| **Total** | | **81** |

**Supplementary table S2**. Body mass estimates and sources for *Leopardus pardinoides* prey species detected from DNA metabarcoding of scat samples.

| Presumed species | Common name | mass (g) | Source | notes |
|---|---|---|---|---|
| Craugastor aenigmaticus | Montane Dirt Frog | 1.5 | citation | |
| Craugastor Sp. | Craugastor frog | 1.5 | citation | |
| Colibri thalassinus | Mexican violetear hummingbird | 5.9 | citation | |
| Microchera albocoronata | Snowcap hummingbird | 2.7 | citation | |
| Panterpe insignis | Fiery-throated hummingbird | 5.7 | citation | |
| Selasphorus scintilla | Scintillant hummingbird | 2.2 | citation | |
| Patagioenas fasciata | Band-tailed pigeon | 366.3 | citation | |
| Zentrygon costaricensis | Buff-fronted Quail-Dove | 200 | VertNet + GBIF | |
| Chamaepetes unicolor | Black Guan | 1135 | citation | |
| Odontophorus guttatus | Spotted Wood-Quail | 303.8 | citation | |
| Basileuterus melanogenys | Black-cheeked Warbler | 11.8 | citation | |
| Cardellina pusilla | Wilson's warbler | 7.5 | VertNet + GBIF | |
| Oreothlypis gutturalis | Flame-throated Warbler | 10.5 | VertNet + GBIF | |
| Atlapetes albinucha | White-naped brushfinch | 33.84 | citation | |
| Pselliophorus luteoviridis | Yellow-green brushfinch | 30 | VertNet + GBIF | |
| Phainoptila melanoxantha | Black-and-yellow Silky-flycatcher | 56 | citation | |
| Scytalopus argentifrons | Silvery-fronted Tapaculo | 17 | citation | |
| Diglossa baritula | Cinnamon-bellied flowerpiercer | 9.2 | citation | |
| Catharus gracilirostris | Black-billed Nightingale-Thrush | 21 | citation | |
| Myadestes melanops | Black-faced Solitaire | 32.1 | citation | |
| Unknown Turdus sp. | Thrush | 71.5 | citation | Median for Turdus |
| Turdus plebejus | Mountain Thrush | 86.33 | citation | |
| Zeledonia coronata | Wrenthrush | 21 | Citation | |
| Pyrrhura hoffmanni | Sulphur-winged parakeet | 82.2 | Citation | |
| Unknown Marmosa Sp. | Woolly mouse opossum | 46.2 | Citation | Median for marmosa |
| Cryptotis gracilis | Talamancan small-eared shrew | 7 | citation | |
| Cryptotis nigrescens | Blackish small-eared Shrew | 7.65 | citation | |
| Unknown Cryptotis Sp. | Small eared shrew | 7.72 | citation | Median for Cryptotis |
| Sylvilagus brasiliensis | Common tapeti cottontail | 950 | citation | |
| Unknown Sylvilagus Sp. | Cottontail rabbit | 1343.4 | citation | Sylvilagus dicei used as is likley species |
| Handleyomys alfaroi | Alfaro's Rice Rat | 26 | citation | |

| | | | | |
|---|---|---|---|---|
| Unknown Handleyomys Sp. | Rice rat | 49.5 | citation | Median for Handleyomys |
| Nephelomys albigularis | Tomes's Rice Rat | 66 | VertNet + GBIF | |
| Unknown Nephelomys Sp. | Rice rat | 60 | VertNet + GBIF | |
| Unknown Nyctomys Sp. | Vesper rat | 49.4 | citation | Nyctomys sumichrasti used |
| Unknown Nyctomys Sp. | Vesper rat | 49.4 | citation | Nyctomys sumichrasti used |
| Oligoryzomys fulvescens | Fulvous pygmy rice rat | 25 | citation | |
| Peromyscus nudipes | Talamancan Deermouse | 24 | VertNet | |
| Reithrodontomys creper | Chiriqui harvest mouse | 22.8 | citation | |
| Reithrodontomys sumichrasti | Sumichrast's harvest mouse | 19 | citation | |
| Reithrodontomys cherrii | Costa Rican Harvest mouse | 14 | VertNet | |
| Scotinomys teguina | Alston's singing mouse | 11.25 | citation | |
| Scotinomys xerampelinus | Chiriqui singing mouse | 15 | citation | |
| Unknown Scotinomys Sp. | Singing mouse | 13.1 | citation | Median for Scotinomys |
| Tylomys watsoni | Watson's climbing rat | 240 | citation | |
| Orthogeomys cavator | Chiriqui pocket gopher | 650 | citation | |
| Heteromys oresterus | Mountain spiny pocket mouse | 74.8 | citation | |
| Unknown Heteromys Sp. | Pocket mouse | 73.65 | citation | |
| Sciurus granatensis | Red-tailed squirrel | 250 | citation | |
| Abronia monticola | Montane alligator lizard | 12.6 | citation | |
| Anolis pachypus | Cloud forest Anole | 3 | citation | |
| Sceloporus malachiticus | Emerald Swift Spiny Lizard | 20 | citation | |
| Bothriechis nubestris | Talamancan palm-pitviper | 230 | citation | |
| Median | | 25 | | |

**Supplementary Table S3.** Arthropod species detected from DNA metabarcoding of *Leopardus pardinoides* scat samples from BOLDigger3 taxonomic assignment.

| Class | order | family | genus | species | % identity |
|---|---|---|---|---|---|
| Arachnida | Mesostigmata | Ascidae | Asca | | 95.207668 |
| Arachnida | Mesostigmata | Phytoseiidae | Typhlodromus | | 100 |
| Arachnida | Sarcoptiformes | Acaridae | Tyrophagus | Tyrophagus putrescentiae | 100 |
| Arachnida | Sarcoptiformes | Scheloribatidae | | | 100 |
| Arachnida | Trombidiformes | Anystidae | | | 100 |
| Arachnida | Trombidiformes | Demodicidae | Demodex | Demodex folliculorum | 99.038462 |
| Arachnida | Trombidiformes | Eriophyidae | Aceria | Aceria tosichella | 99.680511 |
| Arachnida | Trombidiformes | Eupodidae | | | 94.285714 |
| Collembola | Entomobryomorpha | Isotomidae | Desoria | Desoria trispinata | 100 |
| Collembola | Entomobryomorpha | Isotomidae | Desoria | | 95.527157 |
| Collembola | Entomobryomorpha | Isotomidae | Parisotoma | | 100 |
| Copepoda | Cyclopoida | Cyclopidae | Macrocyclops | Macrocyclops distinctus | 99.361022 |
| Insecta | Coleoptera | Cerambycidae | Vadonia | | 95.238095 |
| Insecta | Coleoptera | Cerambycidae | Vadonia | | 95.238095 |
| Insecta | Coleoptera | Cerambycidae | Vadonia | | 95.238095 |
| Insecta | Coleoptera | Cerambycidae | | | 94.047619 |
| Insecta | Coleoptera | Chrysomelidae | | | 96.784566 |
| Insecta | Coleoptera | Corylophidae | | | 99.680511 |
| Insecta | Coleoptera | Elateridae | | | 96.761134 |
| Insecta | Coleoptera | Leiodidae | | | 100 |
| Insecta | Coleoptera | Melyridae | | | 97.763578 |
| Insecta | Coleoptera | Scarabaeidae | Anomala | Anomala subridens | 99.361022 |
| Insecta | Coleoptera | Scarabaeidae | Deltochilum | Deltochilum burmeisteri | 98.722045 |
| Insecta | Coleoptera | Scarabaeidae | Hoplia | Hoplia bilineata | 98.29932 |
| Insecta | Coleoptera | Scarabaeidae | | | 94.444444 |
| Insecta | Coleoptera | Staphylinidae | | | 96.774194 |
| Insecta | Coleoptera | Staphylinidae | | | 96.108949 |
| Insecta | Diptera | Anthomyiidae | Lasiomma | | 100 |
| Insecta | Diptera | Anthomyiidae | | | 99.176955 |
| Insecta | Diptera | Calliphoridae | Calliphora | Calliphora triseta | 100 |
| Insecta | Diptera | Calliphoridae | Compsomyiops | Compsomyiops verena | 100 |
| Insecta | Diptera | Calliphoridae | Lucilia | Lucilia purpurascens | 99.680511 |
| Insecta | Diptera | Cecidomyiidae | Dasineura | | 99.21875 |
| Insecta | Diptera | Cecidomyiidae | | | 98.402556 |
| Insecta | Diptera | Chironomidae | Orthocladius | Orthocladius oblidens | 100 |
| Insecta | Diptera | Chironomidae | | | 100 |
| Insecta | Diptera | Chloropidae | Biorbitella | | 100 |
| Insecta | Diptera | Chloropidae | Malloewia | Malloewia aequa | 98.402556 |
| Insecta | Diptera | Chloropidae | | | 99.680511 |
| Insecta | Diptera | Drosophilidae | Drosophila | Drosophila affinis | 97.107438 |

| | | | | | |
|---|---|---|---|---|---|
| Insecta | Diptera | Drosophilidae | | | 100 |
| Insecta | Diptera | Lauxaniidae | | | 99.678457 |
| Insecta | Diptera | Limoniidae | | | 99.041534 |
| Insecta | Diptera | Muscidae | | | 100 |
| Insecta | Diptera | Phoridae | | | 100 |
| Insecta | Diptera | Psychodidae | | | 100 |
| Insecta | Diptera | Sarcophagidae | | | 94.650206 |
| Insecta | Diptera | Sciaridae | Bradysia | | 100 |
| Insecta | Diptera | Sciaridae | | | 100 |
| Insecta | Diptera | Sphaeroceridae | Sclerocoelus | Sclerocoelus nitidistylus | 99.361022 |
| Insecta | Diptera | Sphaeroceridae | | | 100 |
| Insecta | Diptera | Syrphidae | Ocyptamus | | 99.680511 |
| Insecta | Diptera | Tachinidae | Parepalpus | Parepalpus labeosus | 97.75641 |
| Insecta | Diptera | Tachinidae | | | 94.193548 |
| Insecta | Diptera | Tipulidae | Tipula | Tipula balloui | 99.680511 |
| Insecta | Diptera | Tipulidae | | | 100 |
| Insecta | Diptera | | | | 98.634812 |
| Insecta | Hemiptera | Aphalaridae | | | 100 |
| Insecta | Hemiptera | Cicadellidae | | | 97.124601 |
| Insecta | Hemiptera | Miridae | Phytocoris | | 100 |
| Insecta | Hemiptera | Miridae | | | 94.771242 |
| Insecta | Hemiptera | Rhyparochromidae | Ozophora | Ozophora picturata | 97.328244 |
| Insecta | Hemiptera | | | | 100 |
| Insecta | Hymenoptera | Chrysididae | | | 94.53125 |
| Insecta | Hymenoptera | Formicidae | Labidus | | 98.722045 |
| Insecta | Lepidoptera | Erebidae | Hypena | | 97.269625 |
| Insecta | Lepidoptera | Nepticulidae | Enteucha | | 99.680511 |
| Insecta | Lepidoptera | Noctuidae | Eriopyga | | 99.361022 |
| Insecta | Lepidoptera | Noctuidae | | | 94.244604 |
| Insecta | Lepidoptera | Tortricidae | Argyrotaenia | | 98.402556 |
| Insecta | Psocodea | Liposcelididae | Liposcelis | Liposcelis brunnea | 98.402556 |
| Insecta | Siphonaptera | Pulicidae | Ctenocephalides | Ctenocephalides felis | 98.717949 |
| Insecta | Thysanoptera | Thripidae | Frankliniella | | 100 |
| Malacostraca | Isopoda | | | | 94.888179 |
